# gEVAL – A web based browser for evaluating genome assemblies

**DOI:** 10.1101/038638

**Authors:** William Chow, Kim Brugger, Mario Caccamo, Ian Sealy, James Torrance, Kerstin Howe

## Abstract

**Motivation:** For most research approaches, genome analyses are dependent on the existence of a high quality genome reference assembly. However, the local accuracy of an assembly remains difficult to assess and improve. The gEVAL browser allows the user to interrogate an assembly in any region of the genome by comparing it to different datasets and evaluating the concordance. These analyses include: a wide variety of sequence alignments, comparative analyses of multiple genome assemblies, and consistency with optical and other physical maps. gEVAL highlights allelic variations, regions of low complexity, abnormal coverage, and potential sequence and assembly errors, and offers strategies for improvement. While gEVAL focuses primarily on sequence integrity, it can also display arbitrary annotation including Ensembl or TrackHub sources. We provide gEVAL web sites for many human, mouse, zebrafish and chicken assemblies to support the Genome Reference Consortium, and gEVAL is also downloadable to enable its use for any organism and assembly.

**Availability:** Web Browser: http://geval.sanger.ac.uk, Plugin: http://wchow.github.io/wtsi-geval-plugin.

**Supplementary information:** Supplementary data are available at *Bioinformatics* online.

## 1 Introduction

Reference genomes are the foundation for genomic biology. As more and more de-novo sequencing projects are being conducted, and more draft genomes released, the continued challenge is to create sufficiently complete and correct assemblies that can be confidently used as references by the research community. Although it has been over a decade since it was announced that the Human genome was completed, this statement referred to the then achievable quality of sequence resolution. There remained errors and lack of sequence coverage for important regions, for example many areas encoding multi-gene families (IHGSC, 2004, Horton *et al*., 2008). Since then, genome assemblies of varying quality have been produced for many more species and individuals, often resulting in annotation errors with negative influence on their research use (Denton *et al*., 2014). Groups such as the Genome Reference Consortium (GRC, genomereference.org) lead efforts to curate and resolve genome assembly issues, and maintain the best quality genome references possible for key research species (Church *et al*., 2011).

Depending on the project, groups that tackle creating a genome reference may have at their disposal multiple alternative assemblies together with resources such as clone libraries, collections of short reads, cDNA sequences, RNAseq data, physical maps, optical maps, or genetic markers. These datasets help in repairing and reorganizing genomic regions or to create a strategy to infuse new sequence into a draft assembly. In parallel, research biologists who observe discrepancies between their data and the genome reference would benefit from being able to assess evidence supporting the reference in the region.

Current public genome browsers include many of these types of datasets in their analysis repertoire, but usually focus on gene annotation (Cunningham *et al*., 2014, Rosenbloom *et al*., 2014). They also represent major assembly builds released every few years and neither reflect uncertainties in the sequence itself, nor more recent improvements. Here we introduce the gEVAL Browser project, a collection of software and frequently updated databases for key species that takes a tiling path as the backbone, conducts analyses using new sources of data and regularly releases the results in a web interface for users to evaluate sequence integrity and create strategies for sequence management.

## 2 Overview

The web based gEVAL browser and the underlying back end databases use the Ensembl project as a framework to build on (Birney *et al*., 2004). Supporting the GRC, it features the current, previous and in-progress official reference genomes of human, mouse, chicken and zebrafish while also hosting assemblies of these species not found in other public genome browsers including those curated by the Genome in a Bottle Consortium (Zook *et al*., 2015) (Supplementary Table 1), and 16 lab and wild derived mouse strains from the international Mouse Genomes Project (Adams *et al*., 2015). Other species present but less frequently updated include pig (*Sus scrofa*), rat (*Rattus norvegicus*) and three helminth species (*Echinococcus multilocularis, Strongyloides ratti, Schistosoma mansoni*) (Supplementary Table 1).

There are chromosomal, regional overview, detailed region and comparative browser views, in addition to lists of anomalies. Individual analysis tracks displaying a wide variety of glyphs can be selected interactively. Common tracks include clone library end sequences (Figure 1A), cDNA alignments (Figure 1C), genetic markers and self-alignments within the genome. Additional tracks available for some assemblies include issues in regions under review by the GRC, exact clone or sequence component placements and quality of overlaps between neighbouring assembly components (Figure 1B). The glyphs are coloured to indicate discrepancies and offer pop-up menus that reveal further information and navigational options. All tracks incorporate navigational links to access adjacent problematic regions, allowing the user to “walk” along the genome from one feature/issue to the next.

**Figure 1:**
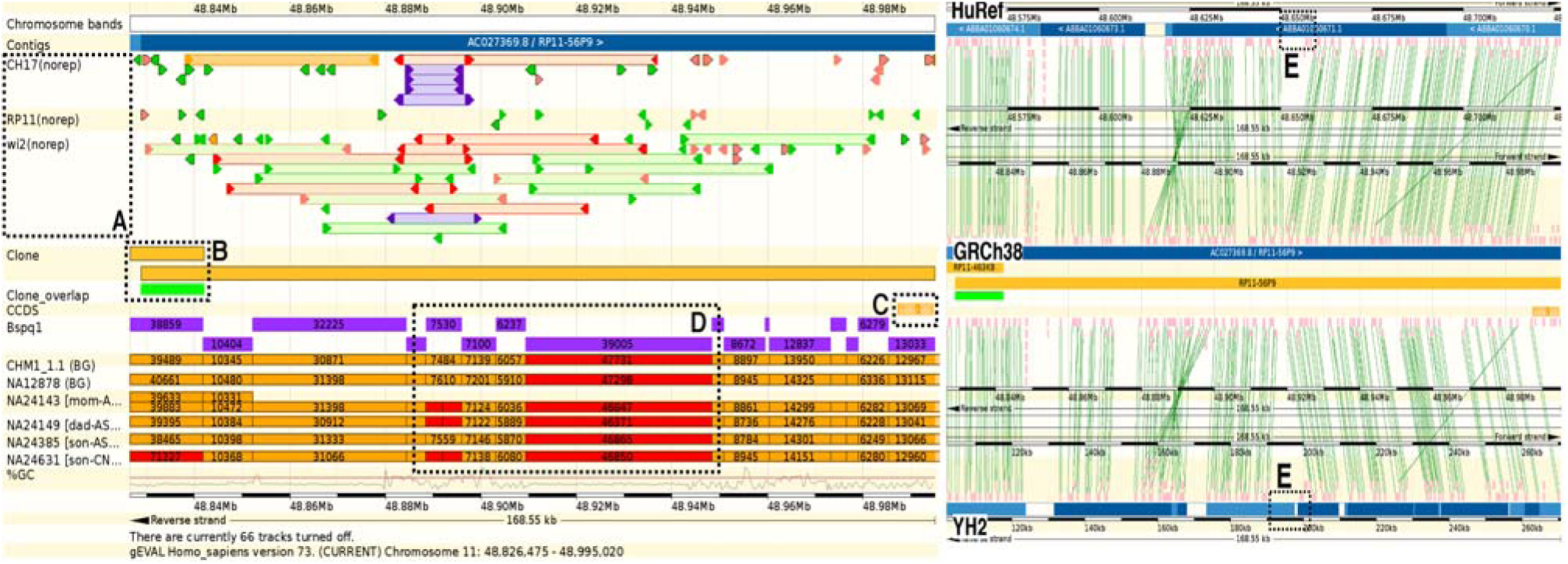
gEVAL analysis of a region on GRCh38 Chromosome 11 with variation and missing sequence. (A) Purple clone end pair mappings indicate same end repeated, while red mappings indicate incorrect orientation of paired ends. (B) Two clone components are used to build this region of the assembly. The green box indicates a reliable overlap region (red would indicate high variation). (C) Orange indicates an incomplete transcript mapping. (D) 6 Single molecule genome maps (orange/red) can be compared to *in silico* digest (purple). Red regions indicate discordance. In this case, a ∽7.5kb block variation is shared between 3 maps and the reference, whilst 3 other maps share two fragments. Furthermore, in the ∽39kb digest block, all 6 maps indicate a size of ∽45-47kb, giving evidence of missing sequence (∽7-8kb). (E) Comparative analysis between HuRef and YH2 assemblies, reveal this missing sequence (dotted box) as well as the region of variation (Supplementary Fig 1).

Comparative genome analysis is invaluable for revealing conserved regions between organisms. Uniquely in gEVAL, comparative analysis focuses on genomic alignments among the different assemblies available for the same species (Figure 1E). This is useful in capturing sequence differences caused by both variation as well as misassembly, and aids the improvement of one assembly with components/guidance from another (Supplementary Figure1).

Unique to gEVAL, it integrates general annotation with long range information produced from recent technologies such as single molecule genome/optical maps (Teague *et al*., 2010, Howe and Wood, 2015) (Supplementary Table 2). These maps prove invaluable for scaffolding assemblies but are also useful for capturing genome-wide structural variation (Mak *et al*., 2015) (Figure 1D). In addition to presenting evidence, gEVAL can suggest specific sequences from an aligned resource to fill reference gaps, or can suggest inversions or other rearrangements such as expansions or contractions of tandem repeats.

## 3 Conclusion

gEVAL is a web-based browser that allows easy detailed evaluation of genome assemblies through its tools and pre-computed analyses, and suggestion of fixes. It is a key tool used by the GRC for the improvement of the primary vertebrate reference genomes. It is widely used externally for evaluating detailed regions of these genomes in different assembly versions, and also by other reference assembly projects, for example the respective international genome consortiums of pig (Groenen *et al*., 2012, Warr *et al*., 2015), chicken (Schmid *et al*., 2015) and mouse (Adams *et al*., 2015). To enable this use the gEVAL code can be downloaded (see Availability Section) and installed alongside an Ensembl installation for any other genome assembly.

## Acknowledgements

The authors would like to thank Richard Durbin for his support and helpful comments on the manuscript.

